# Optimized liquid and gas phase fractionation increase HLA-peptidome coverage for primary cell and tissue samples

**DOI:** 10.1101/2021.05.25.445487

**Authors:** Susan Klaeger, Annie Apffel, Karl R. Clauser, Siranush Sarkizova, Giacomo Oliveira, Suzanna Rachimi, Phuong M. Le, Anna Tarren, Vipheaviny Chea, Jennifer G. Abelin, David A. Braun, Patrick A. Ott, Hasmik Keshishian, Nir Hacohen, Derin B. Keskin, Catherine J. Wu, Steven A. Carr

## Abstract

Mass spectrometry is the most effective method to directly identify peptides presented on HLA molecules. However, current standard approaches often require many millions of cells for input material to achieve high coverage of the immunopeptidome and are therefore not compatible with the often limited amounts of tissue available from clinical tumor samples. Here, we evaluated microscaled basic reversed-phase fractionation to separate HLA peptide samples off-line followed by ion mobility coupled to LC-MS/MS for analysis. The combination of these two separation methods enabled identification of 20% to 50% more peptides compared to samples analyzed without either prior fractionation or use of ion mobility alone. We demonstrate coverage of HLA immunopeptidomes with up to 8,107 distinct peptides starting with as few as 100 million cells or 150 milligrams of wet weight tumor tissue. This increased sensitivity can improve HLA binding prediction algorithms and enable detection of clinically relevant epitopes such as neoantigens.

## Introduction

Antigen presentation and subsequent recognition by cytotoxic T cells is a central component of the adaptive immune response. Therapies harnessing the power of the immune system are showing great potential in personalized cancer treatment ^1–4^. Because these therapies rely on recognition of tumor-associated antigens (TAA), viral antigens or neoantigens by cytotoxic T cells, discovery and identification of these antigens is crucial to impact patient treatment outcomes. Cell surface antigens presented on highly polymorphic human leukocyte antigen class I (HLA-I) molecules are short peptide sequences that are predominantly 9 amino acids long. Which antigens are presented by an individual’s HLA allele is often predicted computationally or evaluated using biochemical binding assays with a biased set of selected synthetic peptides. Mass spectrometry (MS) enables direct, untargeted identification of thousands of endogenously processed and presented peptide antigens and therefore provides a deeper insight into the immunopeptidome ^5–8^. In particular, MS-based detection of potential tumor antigens can confirm the presentation of specific epitopes by tumor cells, identify new antigens with therapeutic potential or monitor changes in antigen presentation in response to treatment ^9–12^.

At present, the identification of HLA-I eluted peptides by mass spectrometry (immunopeptidomics) is hampered by multiple factors. First, HLA-bound peptides span a diverse repertoire of sequences as each HLA-allele has a unique peptide-binding motif. Second, despite their different sequences, these peptides are very similar to each other in length and amino acid composition. Thus, effective separation using standard LC-MS/MS methods optimized on tryptic proteome digests remains challenging. Third, thousands of peptides are estimated to be presented by each cell, each with relatively low abundance and therefore, a large amount of input material is needed to achieve high sensitivity. Fourth, algorithmic interpretation of HLA-I eluted peptide spectra is much more challenging than for trypsin-derived peptide spectra typical of proteomic studies because: a) the narrow length distribution of the complex HLA-I eluted peptides mixture hampers chromatographic separation leading to more co-isolated precursor ions producing mixed MS/MS spectra, b) charge bearing, basic residues (e.g., Lys, Arg) are not always at the C-terminus and may be absent altogether, leading to substantially altered distributions of fragment ion series, and c) the lack of restricted enzyme specificity inflates the sequence database search space by ∼100x ^13^.

While recent improvements in MS instrumentation have made HLA-I eluted peptide identification more feasible, the detection of promising tumor rejection antigens such as neoantigens remains particularly challenging. The amount of HLA-eluted peptide material derived from small amounts of tumor tissue obtained in a clinical setting is often limited. Moreover, extensive sample handling following enrichment of HLA-I eluted peptides can lead to sample loss. Successful tissue-based approaches have typically utilized billions of cells for analysis ^11^. Previously, we developed and optimized methods for isolation of immunoprecipitated HLA-peptide complexes from as few as 50 million cells and 0.2g of clinical specimens yielding 1,000 - 2,000 peptides per sample (**Figure 1**, left panel) ^7,8,14,15^. For subsequent analysis by liquid chromatography tandem mass spectrometry (LC-MS/MS), three HLA:peptide immunoprecipitations (IPs, 150 ⨯ 10^6^) were pooled, desalted and analyzed in two technical replicate injections. The method has yielded sufficient depth (1,000- 2,000 peptides per sample) for profiling of mono-allelic HLA-expressing cell lines and HLA-bound antigens from patient tumor specimens ^8^. While the depth of coverage obtained using such small amounts of sample was high by historical standards, it has generally been inadequate to detect low frequency peptides (i.e. like neoantigens) consistently. Furthermore, we observed that with each repeat injection, we uncovered more of the peptidome; around 20% of peptides were only detected in one of four injections suggesting that there is far more of the peptidome to be discovered (**Supplemental Figure 1A, B)**.

**Figure 1:**
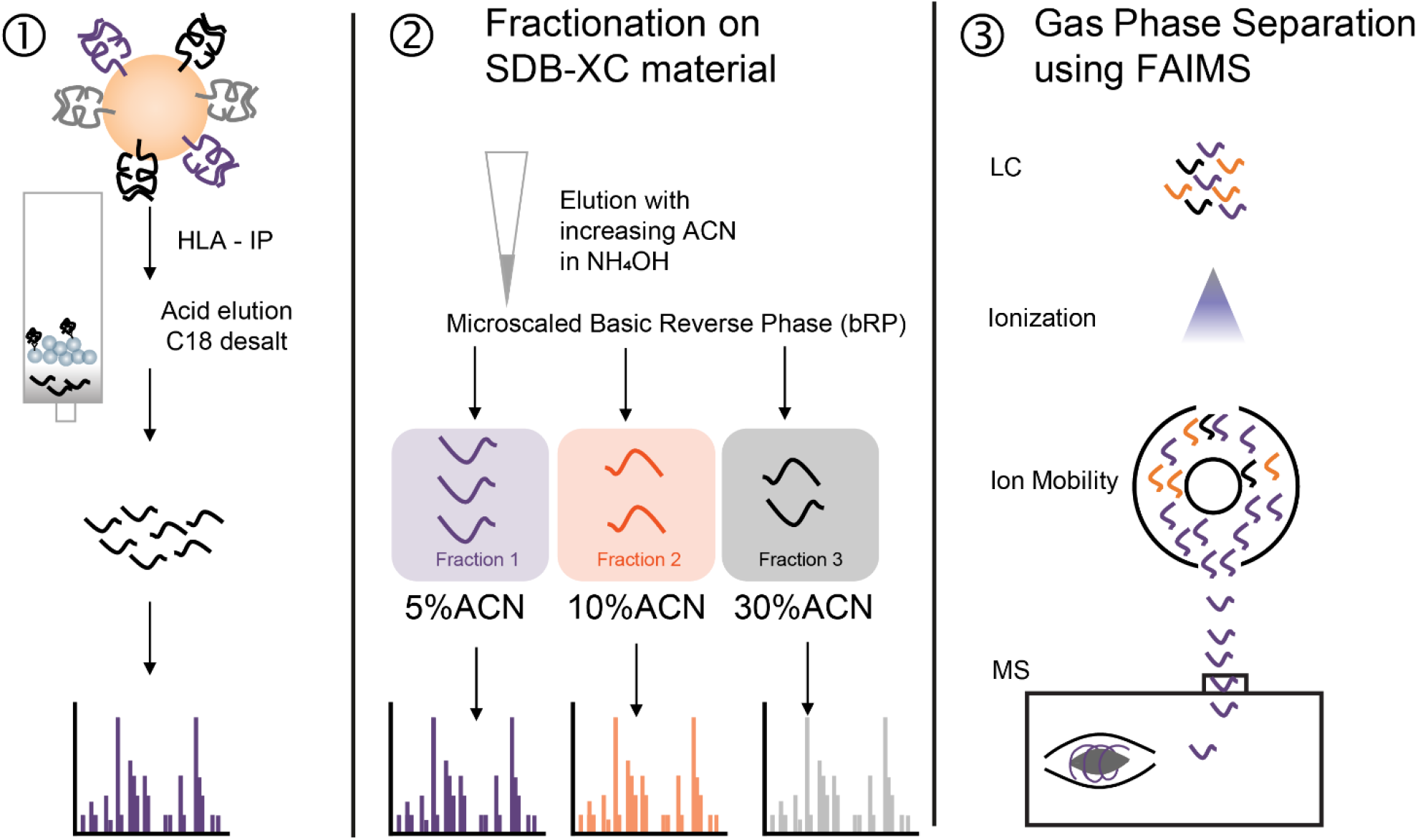
Schematic overview of HLA sample preparation approaches. Prior standard workflow in which HLA-I peptides are acid eluted and desalted before LC-MS/MS analysis with repeat injection. 2) Offline, basic reversed-phase separation of eluted peptides, further separated into three fractions using SDB-XC material using increasing acetonitrile concentrations in ammonium-hydroxide (5%, 10% and 30% ACN in 0.1% NH_4_OH, pH 10). 3) Separation of ions in the gas phase using FAIMS coupled to the mass spectrometer.

Recent improvements in proteome profiling protocols for small sample input amounts and the introduction of ion mobility for gas phase fractionation on-line with the MS system have the potential to improve the depth of HLA-I eluted peptide analysis. Stage-Tip-based basic reversed-phase fractionation using alternative, non-C18 solid phases has already shown benefits for proteomic analysis of low cell numbers after flow cytometry ^16^. In addition, use of high field asymmetric waveform ion mobility spectrometry (FAIMS) on hybrid quadrupole-orbitrap instruments has been demonstrated to increase peptide and protein identifications in single-shot proteomics experiments ^17–19^. The advantage of FAIMS for gas phase separation of low input, posttranslationally modified peptide samples has also been reported ^20^.

Here, we investigated the impact of using offline Stage-Tip fractionation and online gas phase separation by FAIMS to increase the depth of immunopeptidome coverage for cell line and patient tissue samples with relatively low amounts of input HLA-I eluted peptides. We present an optimized HLA-I immunopeptidomics workflow for the analysis of HLA-I eluted peptides derived from multiple tumor types, revealing that a combination of fractionation and FAIMS significantly improves the profiling depth of the HLA-I immunopeptidome. We demonstrate that this new workflow enables detection of neoantigens and further extends large-scale data generation capabilities that can motivate the retraining of epitope prediction algorithms.

## Experimental procedures

### Generation of single-allele cell lines and primary human samples

HLA class I-deficient B721.221 cell lines expressing a single HLA-I allele each were generated as previously described ^8^. Cell lines with stable surface HLA-I expression were generated through selection using 800 μg/ml G418 (Thermo Scientific), followed by enrichment of HLA-I positive cells through up to two serial rounds of fluorescent activated cell sorting (FACS) and isolation using a pan-HLA-I antibody (W6/32; Santa Cruz) on a FACSAria II instrument (BD Biosciences). All human tissues were obtained through Dana-Farber Cancer Institute- or Partners Healthcare-approved Institutional Review Board (IRB) protocols. Conditions for growth and in vitro propagation of MEL and GBM tumor cell lines and of monocyte-derived dendritic cells were described previously ^8^. Human research biopsies were acquired through protocols approved by the Institutional Review Board at DFCI.

For primary tumors and patient-derived cell lines, HLA–peptide complexes were immunoprecipitated from 0.1–0.2 g tissue or up to 50 million cells. Solid tumor samples were dissociated using a tissue homogenizer (Fisher Scientific 150) and HLA complexes were enriched as described below.

### HLA-I peptide enrichment and LC-MS/MS analysis

Soluble lysates from up to 50 million cells and up to 0.2 g from tumor samples were immunoprecipitated with W6/32 antibody (sc-32235, Santa Cruz) as described previously ^8^ (Supplemental Table 1). Iodoacetamide (10 mM) was added to the lysis buffer to alkylate cysteines. HLA-I bound peptides were acid eluted on 1cc 50mg tC18 SepPak Cartridges (WAT054960, Waters) on a vacuum manifold. SepPaks are equilibrated with 200μL MeOH x 2, 100μL 50% ACN/1% FA, 500μL 1% FA x 4, respectively. Beads with HLA-I bound peptides were resuspended in 3% ACN/ 5% FA and transferred to the equilibrated cartridge. Peptides were eluted from HLA-I proteins with 10% acetic acid for 5 minutes twice and then desalted with 4x 500μL 1% FA. Finally, peptides were eluted from the desalt matrix using 250μL 15% ACN/1% FA followed by 250μL 50% ACN/ 1% FA and dried down.

### Basic reversed-phase fractionation for HLA-I peptides

For basic reversed-phase fractionation, a Stage-Tip was prepared using two disks of SDB-XC material (Empore 2240), washed and equilibrated with 100% methanol, followed by 50% ACN/1% FA and 1% FA. Dried peptides were resuspended in 3% formic acid / 5% ACN and loaded onto the Stage-Tip. Peptides were desalted with up to five washes of 1% FA before elution in three fractions with increasing concentrations of ACN (5%, 10%, 30%) in 0.1% (wt/vol) NH_4_OH (28% NH_3_ (wt/vol), 338818, Sigma Aldrich), pH 10. Fractions were dried down in a vacuum concentrator and stored at -80°C until LC-MS/MS data acquisition.

### LC-MS/MS data acquisition

Samples were reconstituted in 3% ACN/ 5% FA prior to LC-MS/MS analysis on a Fusion Lumos or Orbitrap Exploris 480 (Thermo Scientific). A pool of two IPs was injected twice as a technical replicate. Alternatively, when fractionated, the entire fraction was injected once. Peptides were loaded onto an analytical column (25-30cm, 1.9um C18 (Dr. Maisch HPLC GmbH), packed in-house PicoFrit 75 μm inner diameter, 10 μm emitter (New Objective)). Peptides were eluted with a linear gradient (EasyNanoLC 1200, Thermo Fisher Scientific) ranging from 6-30% Solvent B (0.1%FA in 90% ACN) over 84 min, 30-90% B over 9 min and held at 90% B for 5 min at 200 nl/min. MS/MS were acquired in data dependent acquisition. For the Fusion Lumos, settings were described previously ^8^. On the Orbitrap Exploris 480, MS1 spectra were collected until either 100% normalized AGC target or 50 ms maximum injection time were reached. Monoisotopic peak detection was set to “peptide” and “relax restrictions” was enabled; precursor fit filter was set to 50% fit threshold and 1.2 m/z fit window. Dynamic exclusion was 10 seconds. For HLA-I peptides, up to five precursors of charge 1+ between 800-1700 m/z or 10 precursors of charge 2-4+ were subjected to MS/MS acquisition. Precursors were isolated with a 1.1 m/z window, collected with 50% normalized AGC target and 120ms maximum injection time, fragmented at 30% HCD collision energy and acquired at 15,000 resolution.

When FAIMS was attached, spray voltage was increased to 1900 V and FAIMS was set to standard resolution. FAIMS CVs were set to -50 and -70 with a cycle time of 1.5s per FAIMS experiment. Further evaluated settings included CV combinations -40|-60. If CV -20 was used, MS2 spectra were triggered for the 5 most abundant MS1 precursors. MS2 fill time was set to 100ms, and all other parameters were the same as in the no FAIMS setting.

### HLA peptide identification using Spectrum Mill

Mass spectra were interpreted using the Spectrum Mill software package v7.00 (SM) (Broad Institute, proteomics.broadinstitute.org). Tandem MS (MS/MS) spectra were excluded from searching if they did not have a precursor sequence MH+ in the range 600–4,000, had a precursor charge > 5 or had a minimum of <5 detected peaks. Merging of similar spectra with the same precursor m/z acquired in the same chromatographic peak was disabled. Before searches, all MS/MS spectra had to pass the spectral quality filter with a sequence tag length > 1 (that is, minimum of three masses separated by the in-chain masses of two amino acids). The max sequence tag lengths were calculated in the SM Data Extractor module using the ESI-QEXACTIVE-HCD-v2 peak detection parameters. MS/MS spectra were searched against a protein sequence database that contained 98,298 entries, including all UCSC Genome Browser genes with hg19 annotation of the genome and its protein-coding transcripts (63,691 entries), common human virus sequences (30,181 entries) and recurrently mutated proteins observed in tumors from 26 tissues (4,167 entries), as well as 259 common laboratory contaminants including proteins present in cell culture media and immunoprecipitation reagents. Mutation files for 26 tumor tissue types were obtained from the Broad GDAC portal (gdac.broadinstitute.org). Recurrent mutations in the coding region within each of the 26 tumor types (frequency = 3 for stomach adenocarcinoma, uterine corpus endometrial carcinoma; frequency = 5 for adrenocortical carcinoma, pancreatic adenocarcinoma, MEL; frequency = 2 for rest) were included. For tumor samples, patient specific mutations were appended to the database. MS/MS search parameters included: digest: no-enzyme specificity; instrument: ESI-QEXACTIVE-HCD-HLA-v3; fixed modification: cysteinylation of cysteine; variable modifications: carbamidomethylation of cysteine, oxidation of methionine and pyroglutamic acid at peptide N-terminal glutamine; precursor mass tolerance of ±10 ppm; product mass tolerance of ±10 ppm; and a minimum matched peak intensity of 30%.

Peptide spectrum matches (PSMs) for individual spectra were automatically designated as confidently assigned using the Spectrum Mill autovalidation module to apply target-decoy-based FDR estimation at the PSM level of <1% FDR. Score threshold determination also required that peptides had a minimum sequence length of 7, and PSMs had a minimum backbone cleavage score (BCS) of 5. BCS is a peptide sequence coverage metric and the BCS threshold enforces a uniformly higher minimum sequence coverage for each PSM, at least four or five residues of unambiguous sequence. The BCS score is a sum after assigning a 1 or 0 between each pair of adjacent amino acids in the sequence (max score is peptide length - 1). To receive a score, cleavage of the peptide backbone must be supported by the presence of a primary ion type for higher energy collisional dissociation (HCD): b, y or internal ion C terminus (that is, if the internal ion is for the sequence PWN then BCS is credited only for the backbone bond after the N). The BCS metric serves to decrease false positives associated with spectra having fragmentation in a limited portion of the peptide that yields multiple ion types.

PSMs were consolidated to the peptide level to generate lists of confidently observed peptides for each allele using the Spectrum Mill Protein/Peptide summary module’s Peptide-Distinct mode with filtering distinct peptides set to case sensitive. A distinct peptide was the single highest scoring PSM of a peptide detected for each sample. Different modification states observed for a peptide were each reported when containing amino acids configured to allow variable modification; a lowercase letter indicates the variable modification (C-cysteinylated, c-carbamidomethylated).

The precursor isolation purity (PIP) metric calculated by the SM Data Extractor module is the intensity of the precursor ion and its isotopes divided by the total peak intensity in the precursor isolation window used to generate an MS/MS spectrum (combined from the two MS spectra immediately before and after the MS/MS spectrum). PIP < 50% indicates expected contamination by co-fragmented peptides and constitutes the default threshold for disregarding reporter ion quantitation. Identification rate was calculated by dividing the number of identified spectra by the number of filtered spectra that passed initial quality assessment in the SM Data Extractor module.

### Filtering of MS-identified peptides and subsequent data analysis

The list of LC–MS/MS-identified peptides was filtered to remove potential contaminants in the following ways: (1) peptides observed in negative controls runs (blank beads and blank immunoprecipitates); (2) peptides originating from the following species: ‘STRSG’, ‘HEVBR’, ‘ANGIO432’, ‘ANGIO394’, ‘ANGIO785’, ‘ANGIO530’, ‘ACHLY’, ‘PIG’, ‘ANGIO523’, ‘RABIT’, ‘STAAU’, ‘CHICK’, ‘Pierce-iRT’, ‘SOYBN’, ‘ARMRU’ and ‘SHEEP’ as common laboratory contaminants including proteins present in immunoprecipitation reagents (note that ‘BOVINE’ peptides derived from cell culture media were not excluded as they appear to have undergone processing and presentation and exhibit anchor residue motifs consistent with the human peptides observed for each allele); (3) peptides for which both the preceding and C-terminal amino acids were tryptic residues (R or K). Subsequent data analysis was performed in R using in-house scripts and the ggplot and Upset ^21,22^ packages.

### HLA peptide prediction

HLA peptide prediction was performed using HLAthena ^8^. Unless otherwise specified, peptides were assigned to an allele using a percentile rank cutoff ≤ 0.5.

### Non-metric multidimensional scaling (NMDS)

A pairwise relative peptide distance matrix was computed between every pair of 9-mer peptide sequences identified in mono-allelic datasets evaluated in the different approaches. The distance between two peptides is defined as the sum of AA residue dissimilarities at each position along the sequences. Dissimilarity is weighted by the entropy at the corresponding position ^7^. NMDS was used to reduce the dimensionality of the distance matrix so peptides can be visualized as points in 2D.

### Multiple reaction monitoring (MRM) for neoantigen detection

Multiple reaction monitoring (MRM) assay configuration using heavy labeled synthetic peptides was done on TSQ Quantiva triple quadrupole mass spectrometer (Thermo Fisher) coupled with Easy-nLC 1200 ultra-high pressure liquid chromatography (UPLC) system (Thermo Fisher). Skyline Daily Targeted Mass Spec Environment (version 19) was used throughout assay configuration and all data analysis. First, spectral libraries for the peptides were generated on a Fusion Lumos (Thermo Fisher). Spectral libraries were uploaded to Skyline and 5-10 most intense fragment ions (transitions) for each peptide were selected for MRM assay configuration. Next, collision energies (CE) were optimized for all the transitions and peptides by liquid chromatography – multiple reaction monitoring mass spectrometry (LC-MRM/MS) on TSQ Quantiva using Skyline’s CE optimization module. For every transition starting with the instrument specific calculated CE we tested 10 additional CEs (5 below and 5 above the calculated CE) in increments of 2. The list of transitions with varying CE values was exported from Skyline and used for building MRM method in Xcalibur software. Equimolar mixture of peptides at 50fm/ul was analyzed by LC-MRM/MS on Quantiva using this method. Resulting data was analyzed on Skyline which then selected the CE that resulted in the highest peak area for each transition. In the final step of CE optimization MRM data was acquired with optimized CE values for every transition. Using this dataset in Skyline, the best 2-6 transitions for every peptide were selected manually giving highest priority to fragment ions of y-series with mass to charge (m/z) above the precursor. For the peptides where the options were more limited we also included ions of y-series with m/z below the precursor and b-series.

After CE optimization, we analyzed three replicates of 6 IPs pooled with either 20 fmol or 40 fmol heavy peptide spiked in.

Liquid chromatography was performed on a 75um ID picofrit columns packed in-house to a length of 28-30cm with Reprosil C18-AQ 1.9um beads (Dr Maisch GmbH) with solvent A of 0.1% formic acid (FA) / 3% acetonitrile (ACN) and solvent B of 0.1% FA / 90% ACN at 200nL/min flow rate and the same gradient as described above. MS parameters include 1.5 sec cycle time, Q1 and Q3 resolution of 0.4 and 0.7, respectively, RT scheduling window of 10min.

### TCR reconstruction and expression in T cells for reactivity screening

TCRs selected from the Oliveira et al study were cloned and transduced in donor T cells (Oliveira et al., in revision). Briefly, the full-length TCRA and TCRB chains, separated by Furin SGSG P2A linker, were synthesized in the TCRB/TCRA orientation (Integrated DNA Technologies) and cloned into a lentiviral vector under the control of the pEF1α promoter using Gibson assembly (New England Biolabs Inc). For generation of TCRs, full-length TCRA V-J regions were fused to optimized mouse TRA constant chain, and the TCRB V-D-J regions to optimized mouse TRB constant chain to allow preferential pairing of the introduced TCR chains, enhanced TCR surface expression and functionality ^23,24^.

Donor T cells enriched from PBMCs using PanT cell selection kit (Miltenyi) were activated with antiCD3/CD28 dynabeads (Thermo Fisher Scientific) in the presence of 5ng/mL of IL-7 and IL-15 (PeproTech). After 2 days, activated cells were transduced with a lentiviral vector encoding the TCRB-TCRA chains. Briefly, lentiviral particles were generated by transient transfection of the lentiviral packaging Lenti-X 293T cells (Takahara) with the TCR encoding plasmids and packaging plasmids (VSVg and PSPAX2) ^25^ using Transit LT-1 (Mirus). Lentiviral supernatant was harvested 2 days later, for 2 subsequent days, and used to transduce activated T cells. Lentiviral transductions were performed by inoculation of the virus at 2,000 rpm, 37°C for 2 hours, and cells were cultured on viral supernatant for 3 days. Six days after activations the beads were removed from culture and expanded in media enriched with IL7 and IL15. Transduction efficiency was determined by flow cytometric analysis using the anti-mTCRB antibody. Transduced T cells were used at 14 days post-transduction for TCR reactivity tests. Briefly 2.5×10^5^ TCR-transduced T cells were put in contact with the following targets: i) patient-derived melanoma cell lines (0.25×10^5^ cells); ii) patient PBMCs (2.5×10^5^ cells); iii) patient EBV-LCLs (2.5×10^5^ cells/well) alone or pulsed with peptides; iv) medium, as negative control; v) PMA (50 ng/ml, Sigma-Aldrich) and ionomycin (10 μg/ml, Sigma-Aldrich) as positive controls. Peptide-pulsing of target cells was performed by incubating EBV-LCLs in FBS-free medium at a density of 5×10^6^ cells/ml for 2 hours in the presence of individual peptides (10^7^ pg/ml, Genscript). After an overnight incubation, TCR reactivity was measured through detection of CD137 surface expression (PE-anti-human CD137, Biolegend) on CD8+ TCR transduced (mTCRB+) T cells using a Fortessa flow cytometer (BD Biosciences).

## Results

### Approaches for HLA peptidome isolation

In an effort to improve sensitivity for detection of HLA-I eluted peptides, we evaluated the impact of microscaled basic reversed-phase fractionation (bRP, **Figure 1** middle) as well as gas phase separation using FAIMS (**Figure 1** right). The effect of microscaled fractionation and FAIMS on the coverage of the immunopeptidome was evaluated individually and in combination, as described below. For all evaluations, we enriched HLA-I peptide complexes from 100×10^6^ cells (i.e. 2 IPs, each at 50×10^6^) that each expressed a single allele ^7,8^ or from up to 0.2g of primary tumor or tumor-derived cell line material using the pan-HLA class I antibody W6/32 (**Supplemental Table 1**). HLA-bound peptides were isolated using acid elution followed by a standard desalt on C18 SepPaks using a stepped elution of 15% and 50% ACN in 0.1% FA. The tested tumor-derived cell lines originated from melanoma (MEL) ^1^, glioblastoma (GBM) ^2^ or clear cell renal cell carcinoma (RCC) tumor specimens.

### Basic reversed-phase fractionation increases peptidome coverage

Although the value of sample pre-fractionation to increase coverage of the immunopeptidome has been reported by several groups ^26–28^, most studies utilize billions of cells as input for HLA peptide enrichment. We aimed to decrease the amount of input sample required to the more readily obtainable amount of 100×10^6^ cells by testing microscaled fractionation on bRP on Stage-Tips (see **Methods**). We first focused on comparing the yields of HLA-I peptides eluted from two mono-allelic cell lines (B*15:01, B*07:02) fractionated using two different reversed-phase materials, SDB-RPS and SDB-XC, to the use of two technical replicates. In our comparison, SDB-XC more efficiently retained and separated HLA-I eluted peptides than SDB-RPS under basic elution conditions resulting in 1,037 (48%) more unique peptide sequences identified than the non-fractionated sample of the same allele (**Supplemental Figure 2A**). Thus, SDB-XC was used for all further evaluations.

**Figure 2:**
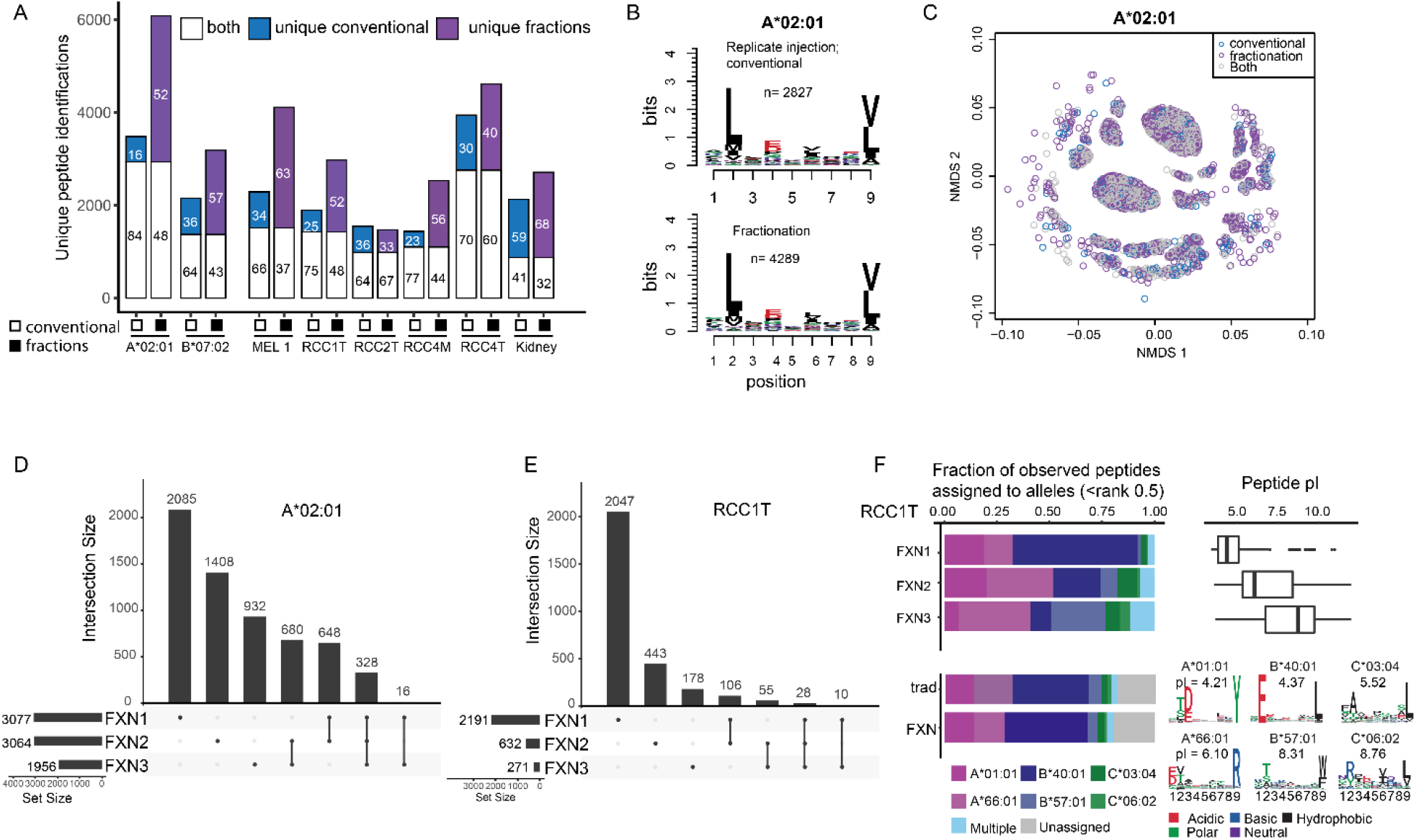
Basic reversed-phase separation increases peptide yield. The number of unique peptides identified by LC-MS/MS obtained without off-line fractionation (empty squares, “conventional” on x-axis) and using fractionation (filled black squares) across mono-and multiallelic samples. Unfilled bars correspond to peptides identified in both approaches; blue filled bars are peptides unique to unfractionated analyses while purple filled bars are peptides unique to fractionated samples. Numbers in bars indicate the percent of total peptides per approach. **B**) Motif of 9-mer peptides identified in the unfractionated (top) and fractionated (bottom) acquisition of the mono-allelic HLA-A*02:01 sample. **C**) Non-metric multidimensional scaling (NMDS) plot (see **Methods**) showing clusters of peptides identified in both (grey), unfractionated (blue) and fractionated (purple) samples of A*02:01 cells. **D**) UpSet plot showing peptide spectrum matches (PSMs) identified in the three fractions of the A*02:01 peptidome. Horizontal bars at bottom left indicate the number of total PSMs per fraction; dots and lines under the vertical bars identify peptides identified in only one (single dot) or multiple fractions (dots with lines). **E**) Same as d) but for RCC1T. **F**) Allele assignment of peptides identified in the individual fractions of RCC1T (top left, fraction of assigned peptides at *HLAthena* rank <0.5) as well as in conventional and fractionated RCC1T (bottom left, fraction of peptides at HLAthena rank <0.5). Top right boxplot shows peptide pI of peptides per fraction, allele motifs and median pI of all peptides assigned to each allele are depicted at the bottom.

We observed a 25-50% gain in peptide identifications for mono-allelic cell lines as well as multi-allelic samples by Stage-Tip fractionation using the SDB-XC bRP material compared to our conventional approach (**Figure 2A**). For example, peptides derived from 100×10^6^ B721.221 cells only expressing HLA-A*02:01 injected in two technical replicates yielded 3,483 unique peptides on the Orbitrap Exploris 480 (Thermo Fisher Scientific). Stage-Tip fractionation of the same amount of HLA-I eluted peptides nearly doubled the yield to 6,084 unique peptides, of which 2,936 peptides were identified by both workflows. The overall peptide yields for a melanoma tumor-derived cell line (MEL1) as well as tissue samples of renal cell cancer (RCC1, RCC4M) increased by 30% relative to our prior approach (**Figure 2A**).

To confirm that the peptides newly identified using bRP fractionation were *bona fide* HLA-I binding peptides, we evaluated how well their sequences matched known HLA-I binding motifs and submotifs. For the A*02:01 eluted peptides, the motifs of technical replicate injection and fractionation revealed the same anchor residues (**Figure 2B**), and the submotifs from these two approaches overlapped when clustered based on their amino acid sequences by non-metric multidimensional scaling (NMDS, **Figure 2C, Methods**) suggesting that the additional peptide identifications obtained from fractionation were likely to bind to HLA-A*02:01 without introducing novel peptide motifs.

To better understand why we obtained a distinct subset of HLA-I eluted peptides using the fractionation approach, we evaluated the peptides’ physiochemical properties in our fractionated dataset. The number of unique peptide identifications per fraction indicated a reasonable separation of peptides based on hydrophobicity and retention on the Stage-Tip in basic conditions. For the HLA-A*02:01 mono-allelic line, 3,077 peptides were identified in the first fraction, 3,064 peptides in the second fraction, and 1,956 peptides in the third fraction. Because peptides were still quite similar in overall properties, 648 and 680 peptides eluted in both fractions 1 and 2 or in fractions 2 and 3, respectively, and 328 peptides were identified in all three fractions (**Figure 2D**). Similar trends were observed for the mono-allelic B*07:02 line (**Supplemental Figure 2D**).

The gain in identifications upon bRP fractionation was even more pronounced for multi-allelic samples, particularly when their peptidomes were derived from binding motifs with a diverse set of physiochemical characteristics (hydrophobicity and isoelectric point -pI). For example, the majority of newly identified peptides for the bRP fractionated human renal cancer tissue sample RCC1 were found in fractions 1 and 2 (5% and 10% cutoff; **Figure 2E, Supplemental Figure 2E** for MEL1). Using the HLA peptide presentation prediction algorithm *HLAthena* ^*8*^, each identified peptide was scored against each patient allele and assigned to a specific allele with a rank cutoff < 0.5. This analysis revealed that under bRP conditions at pH10, peptides separated based on their interaction with the mobile and stationary phase dependent on allele binding motif, hydrophobicity and peptide pI, (**Figure 2F, Supplemental Figure 2F** for MEL1). The median pI of identified peptides in fraction 1 across all alleles was 4.5 and 89% (n=1,111) of B*40:01 binders eluted in fraction 1 (median pI = 4.37). Since the binding motif of B*40:01 has a preference for glutamic acid (E) in position 2, E-containing peptides were likely to be net-negatively charged in our basic buffer conditions and thus not well-retained on the SDB-XC bRP stationary phase. Conversely, peptides associated with alleles bearing basic amino acid residues in the anchor positions, A*66:01 and C*06:02, will tend to be positively charged, retained better and contributed more peptides to the later fractions. Similarly, peptides associated with alleles having hydrophobic residues in their anchor positions, such as B*57:01, tended to preferentially contribute to later bRP fractions (**Figure 2F**). Taken together, bRP fractionation separated peptides in an orthogonal manner to the acidic reversed-phase packing material used in the online gradient and yielded significantly more HLA-I eluted peptides identifications in only 6 hours of measurement time compared to replicate technical replicate analyses that take 4 hours.

### HLA peptide identification benefits from separation of ions in the gas phase

Next, we investigated how the separation of ions by their ion mobility in the gas phase can benefit HLA-I eluted peptide analysis. For these studies, we used the recently introduced FAIMSpro interface from Thermo Scientific that fits orbitrap instruments (e.g., Fusion Lumos and Orbitrap Exploris 480). We tested a range of compensation voltages and optimized the acquisition method for HLA-I eluted peptides using two compensation voltages -50 and -70 (CV), triggering MS2 for 1.5 sec at each CV. Each sample set was acquired on the same instrument (either Fusion Lumos or Exploris, Supplemental Table 1) with and without FAIMS and two replicate injections of the same sample amount (100⨯10^6^ cells). The use of FAIMS increased the number of HLA-I eluted peptides identified by an average of 58% across all samples evaluated (**Figure 3A**). For the peptidome acquisition of the mono-allelic B*53:01-expressing cells and the multi-allelic MEL1 cells, FAIMS acquisition doubled the number of peptides identified compared to no FAIMS. In the case of the RCC1 tumor samples, FAIMS did not increase the overall peptide identifications, but rather revealed 655 new HLA-I eluted peptides not identified using the conventional approach. The main peptide motif of identified peptides using FAIMS matched the motif obtained using conventional approaches (**Figure 3B**). They overlapped in their submotif clusters as illustrated by A*02:01 and B*53:01 (**Figure 3C**) and had similar length distributions compared to the traditional injection (**Supplemental Figure 3A**). Altogether, these characterizations indicate that the new identifications were credible HLA-I binders.

**Figure 3:**
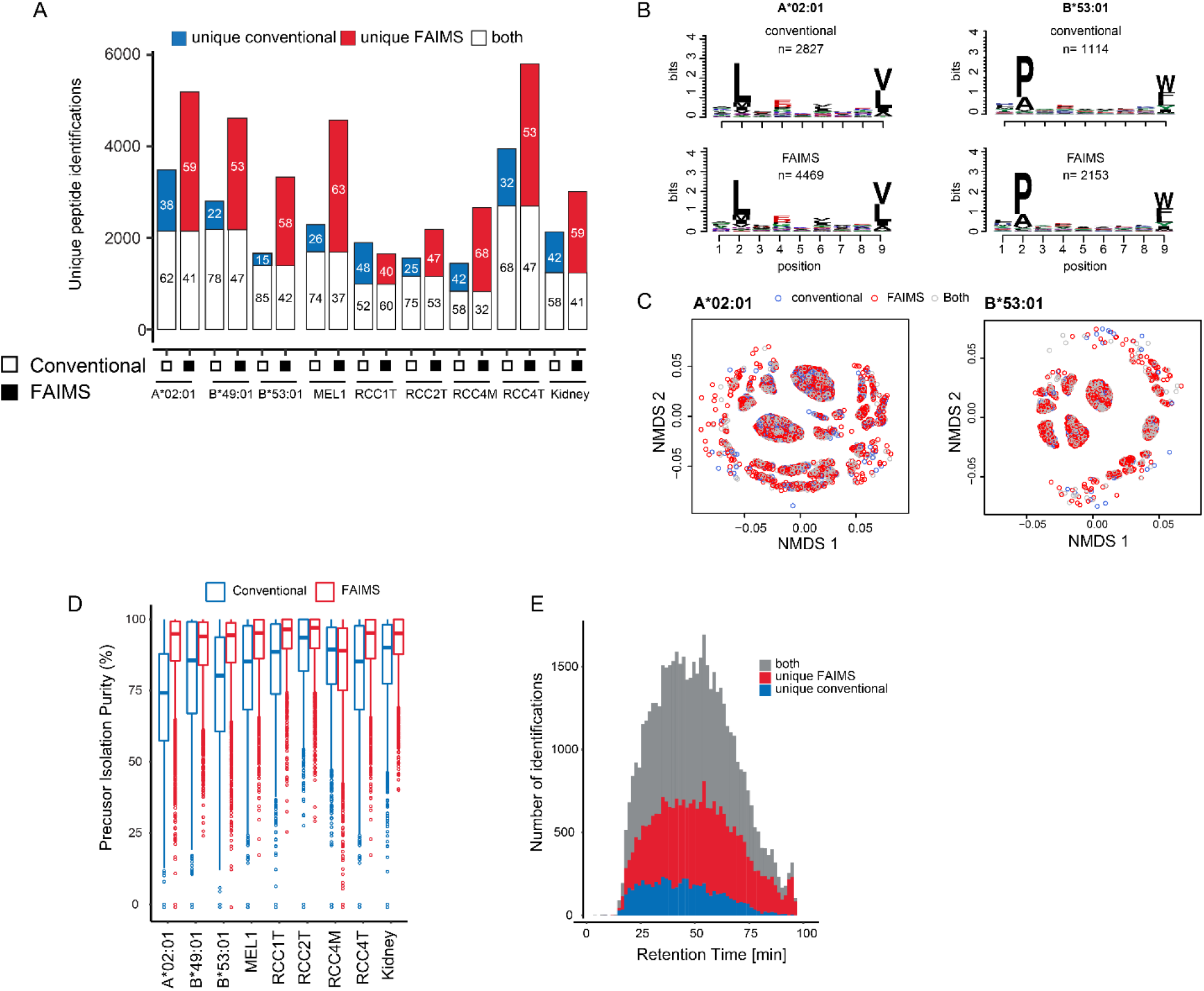
Ion mobility separation increases peptide yield. **A**) The number of HLA-I eluted peptide identifications obtained using FAIMS (filled black squares) or without FAIMS (empty squares) across mono-and multiallelic samples. Unfilled bars correspond to peptides identified in both approaches; blue filled bars are peptides unique to no-FAIMS (conventional) while red filled bars are peptides unique to samples analyzed with FAIMS. Numbers in bars indicate the percent of peptides per approach. **B**) Motif of 9-mer peptides identified in the without (top) and with FAIMS (bottom) acquisition of the mono-allelic HLA-B*53:01 sample on the Orbitrap Exploris. **C**) NMDS plot of peptides identified in both (grey), conventional (blue), and with FAIMS (red) samples of B*53:01. **D**) Precursor isolation purity of peptides in FAIMS (red) and conventional (blue) acquisition. **E**) Number of identifications in all samples across the chromatographic gradient colored by detection in normal acquisition, (no FAIMS, blue), + FAIMS acquisition (red) and both acquisition modes (grey, bin size = 30).

The increased number of identified HLA-I eluted peptides appeared to be related to the overall improved data quality obtained using FAIMS, which facilitated spectral interpretation and identification. FAIMS increased the precursor isolation purity (see **Methods**) from a median between 74-93% to 89-97% (**Figure 3D**). Hence, fewer co-isolated peptide precursor ions competed for one peptide spectrum match (PSM), yielding fewer chimeric spectra and more identifications overall. FAIMS separation facilitated the identification of more precursor ions throughout the whole run but especially increased the yield in the middle of the gradient, when most of the peptides eluted from the column (**Figure 3E**). Total ion chromatograms of individual CV values showed that FAIMS helped to separate ions before they enter the MS (**Supplemental Figure 3B**), and therefore purer MS/MS spectra can be acquired and identified. While FAIMS acquisition with two CVs triggered acquisition of fewer MS/MS spectra than without FAIMS, a higher percentage (identification rate) of the acquired spectra led to an identified peptide sequence in the FAIMS run for all samples (e.g. A*02:01: 30.5% vs 11.5%, **Supplemental Figure 3C**). For samples acquired on the Orbitrap Exploris we observed up to a threefold increase in the percent of identified spectra relative to no FAIMS (**Supplemental Figure 3C**).

Hydrophobic HLA-I eluted peptides lacking basic residues can be observed as singly charged precursor species (0-40%) in the conventional acquisition (**Supplemental Table 2**), but were lost using the two CV settings for FAIMS described above despite allowing for z=1 during precursor selection in the instrument method. To improve the transmission of singly charged peptides, we tested adding an additional CV of -20 which should be better suited to stabilize z=1 precursors ^17^. Peptides derived from mono-allelic cell lines expressing either A*11:01 or B*07:02 were used to evaluate the effect. While A*11:01 features a tryptic-like motif with lysine in the C-terminal position, B*07:02 is representative of a nonpolar peptide pool with a xPxxxxxxL motif. FAIMS experiments were run on peptides from both alleles using CV settings of -50|-70 and CVs of -20|-45|-65. Both methods yielded a similar number of identifications, but while less than 0.1% of peptide spectrum matches (PSMs) were identified with a single charge in A*11:01, CV-20 increased the percentage of singly charged PSMs for B*07:02 to 4% (Supplemental Fig 3D). In another comparison, we evaluated CV combinations -20|-40|-60 and -20|-50|-70 and found that for B*07:02, 6% of singly charged ions led to a peptide identification at CV -20. Since most HLA-I eluted peptides were identified at CV-50 or -60, one of these two CVs should be used for data acquisition in combination with a second CV that stabilizes multiply charged ions. If the sample composition includes alleles with hydrophobic anchor residues known to yield high proportions of singly charged precursors (**Supplemental Table 2**), a third FAIMS experiment at CV -20 and precursor filter for z=1 can be included (**Supplemental Figure 3E**).

In summary, these studies demonstrate that use of the FAIMSpro interface leads to the identification of novel, valid peptides due to improved precursor separation/selection and generation of purer MS/MS spectra.

### Each individual workflow variation can add a distinct set of peptides

Given the increases in depth of HLA-I peptidome coverage obtained using optimized, microscaled bRP fractionation and ion mobility independently, we next evaluated the effect of combining these two approaches. We found that for 4 of 6 samples tested, combining fractionation plus FAIMS yielded more peptides than fractionation alone (**Figure 4A**). However, this was not universally the case, and even in cases where it was true, each method employed also often revealed distinct subsets of peptides.

**Figure 4:**
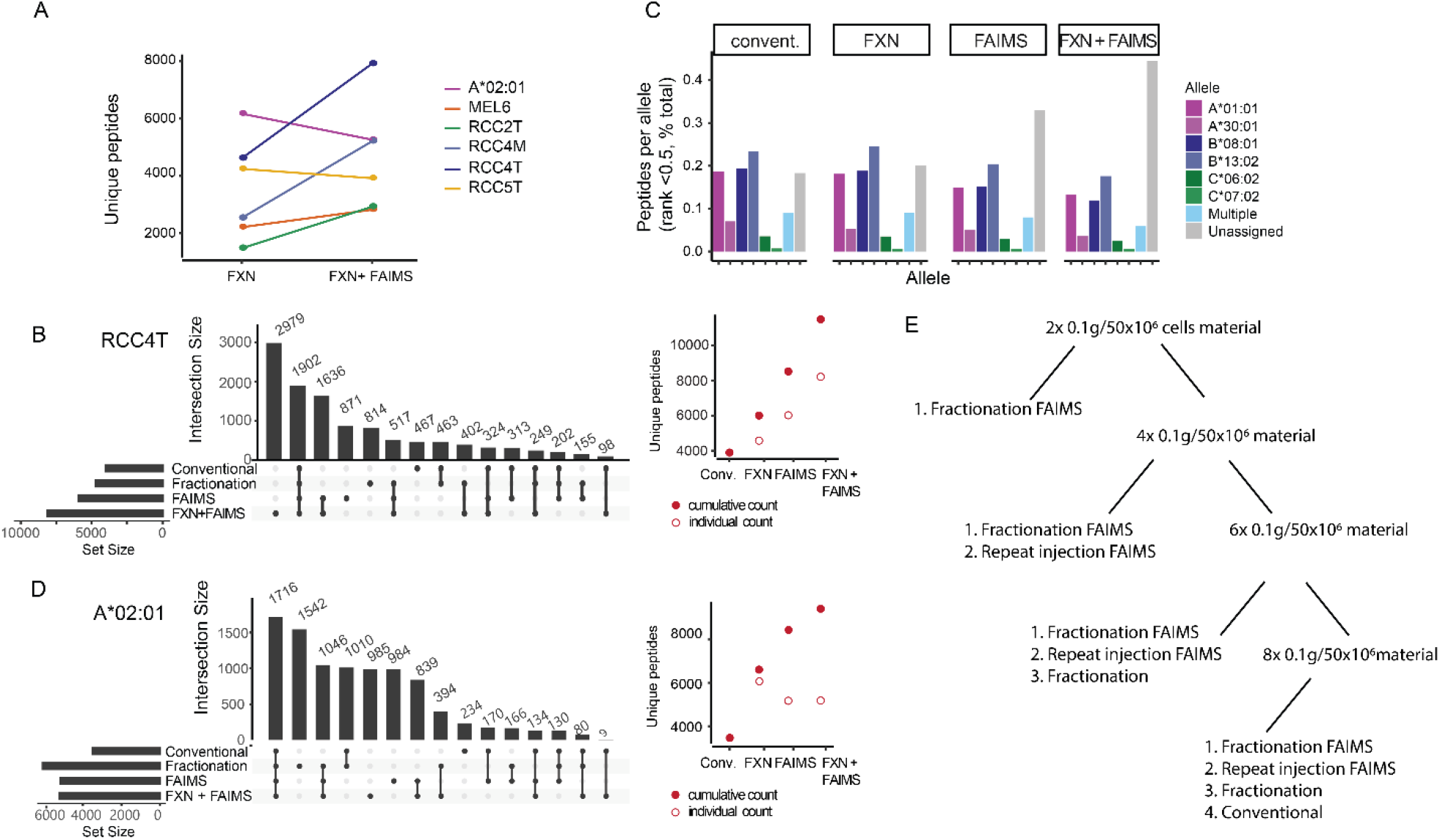
Combination of fractionation and FAIMS for greater coverage of the immunopeptidome. **A**) Number of unique 8-11mer identifications in fractionated samples acquired with and without FAIMS. **B**) Peptides identified across all modes of acquisition of RCC4T. Horizontal bars at bottom left indicate the number of total unique peptides per acquisition; dots and lines under the vertical bars identify peptides identified in only one (single dot) or multiple acquisitions (dots with lines). Each acquisition (empty circle) adds unique peptide identifications to the total cumulative peptidome (filled circle). **C**) Allele assignment of peptides identified in RCC4T (HLAthena, rank < 0.5) across four different acquisitions. **D**) Same as b) but for HLA-A*02:01 sample. **E**) Decision tree for recommended sample preparation and data acquisition strategies based on input available.

In the case of the multi-allelic patient sample RCC4T, off-line Stage-Tip fractionation plus FAIMS added the most unique peptides found in that experiment, followed by using FAIMS alone (**Figure 4B**). In total, 11,392 8-11mers were identified for this patient sample using only 8 IPs originating from 0.1g of tumor material each. Peptides were assigned to the allele to which they were most likely bound (A*01:01, A*30:01, B*08:01, B*13:02, C*06:02, C*07:02) using *HLAthena* ^8^ and a rank cutoff of 0.5. The relative contribution per allele to the total peptidome was similar across different acquisition schemes with the majority of peptides presented on A*01:01, B*08:01 and B*13:02 (**Figure 4C**).

A relatively high percentage of the HLA-I eluted peptides in the FAIMS (37%) and fractionated+FAIMS (52%) samples of RCC4T could not be assigned to an allele at rank <0.5. Half of those peptides were weak binders with rank <2 (**Supplemental Figure 4A**). Closer investigation of the motif of the non-binders at rank <2 suggested potential additional binders for HLA A*30:01 and HLA B*13:02 (**Supplemental Figure 4B**). These alleles have a low PPV in HLAthena ^8^ and hence, binding prediction models can still be improved once more peptides are identified (see below).

For the RCC2 tumor sample, a total of 3,936 HLA-I peptides were detected across all four experiments, while only 773 peptides were commonly identified in all acquisitions (**Supplemental Figure 4C**). The lower yield compared to RCC4T is likely due to plentiful necrotic tissue and diminished HLA peptide presentation. Fractionation in combination with FAIMS alone identified 75% of all peptides and had similar contribution per allele to the total peptidome as the other acquisition schemes.

Applying all variations of the four workflows to a common pool of the mono-allelic HLA-A*02:01 sample yielded a total of 9,439 unique peptides (**Figure 4D**) with varying peptide identifications in each acquisition mode. Of those, only 1,716 were identified across all four conditions. Acquisitions without FAIMS (conventional and fractionation) added 2,786 (1542+1010+234) peptides, while 2,808 additional identifications were found only in acquisitions that used FAIMS (985+984+839). HLA-I eluted peptide yield was greatest for a fractionated sample acquired without FAIMS. In this particular case, the gap in performance between FAIMS and no FAIMS on a QC standard sample (10ng Jurkat proteome tryptic digest) was less pronounced than usual. Hence, maintaining optimal instrument performance is an important consideration.

As each method identified another distinct set of peptides presented on HLA-I, we suggest that greater coverage of the immunopeptidome can be achieved by applying multiple peptide separation approaches rather than performing repeated injection, if sufficient input material is available. Based on our results, when aiming at achieving greatest coverage, we propose a decision tree to guide through sample preparation and acquisition methods (**Figure 4E**). For most tumor samples evaluated in this study, fractionation in combination with FAIMS acquisition resulted in the highest yield of peptides from one sample and enabled neoantigen identifications (see next paragraphs) and hence it was chosen as the preferred sample preparation and data collection method. However, depending on the amount of sample available, FAIMS acquisition without fractionation added another larger set of distinct peptides and can therefore be chosen as an alternative to technical replicates. This was followed by fractionation without FAIMS. Although our conventional approach was outperformed by all other strategies, it still revealed additional peptides not observed in any of the other acquisitions.

### Increased coverage can enable the detection of clinically relevant neoantigens

Increased numbers of identified peptides from lower amounts of tumor sample input can also increase the likelihood of detecting clinically relevant epitopes such as neoantigens. Using either fractionation, FAIMS or both, we profiled different samples from primary tumor as well as from tumor-derived cell lines. The underlying tumor types had different mutational burdens, with MEL harboring more mutations compared to GBM or RCC.

In two melanoma samples, MEL1 and MEL6, we were able to detect 4 and 2 neoantigens respectively, starting with only 100 ⨯10^6^ cells or 0.2g of input per sample using fractionation and FAIMS acquisition (**Figure 5A**). These neoantigens were missed in analyses of these samples using the conventional workflow as well as prior runs of the same tumor derived cell lines ^8^. For GBM and RCC, no neoantigens were detected using these input amounts.

**Figure 5:**
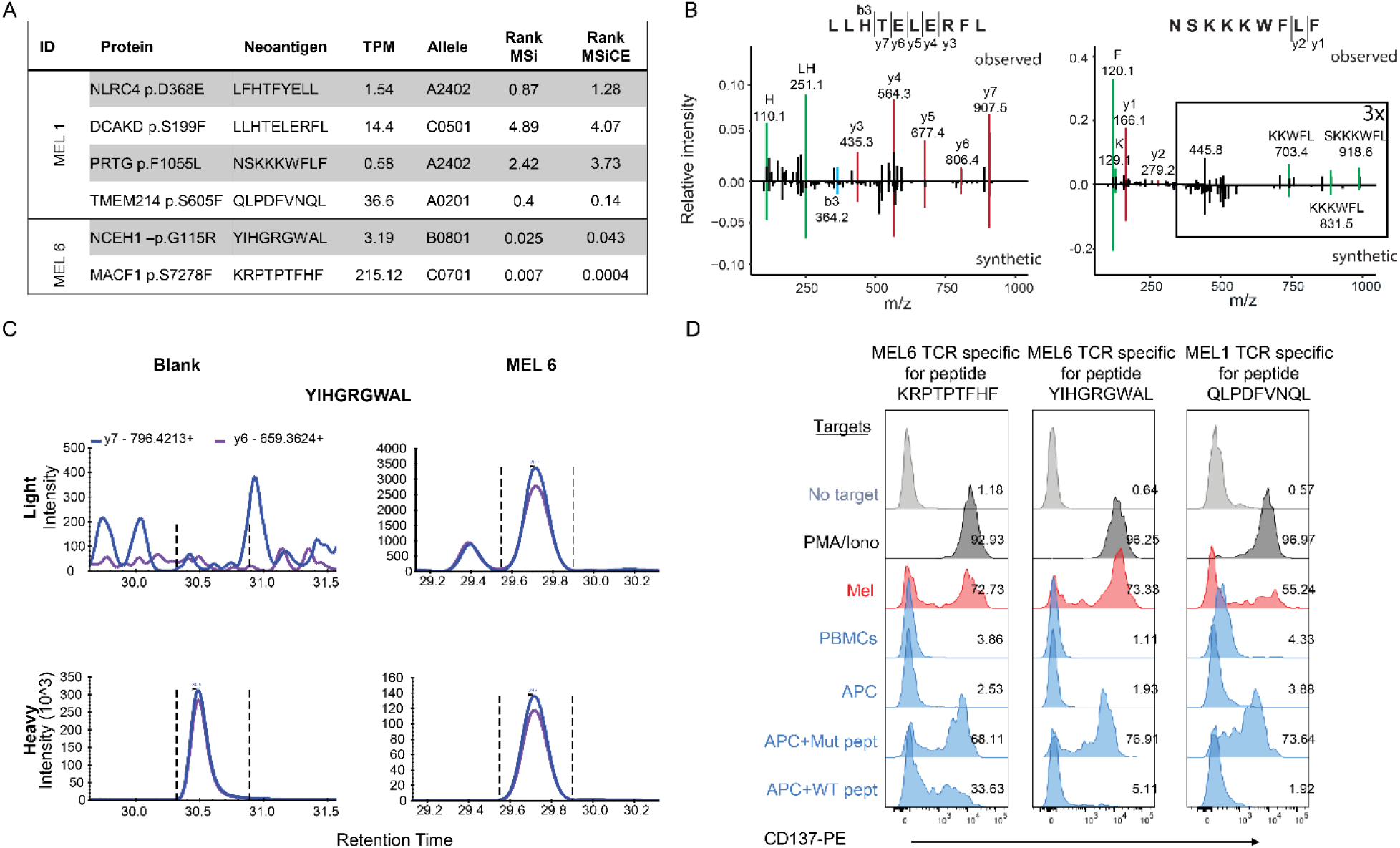
Optimized data acquisition strategies facilitate neoantigen discovery. **A**) Neoantigens identified in MEL 1 and MEL 6 acquired with fractionation +/-FAIMS, their protein of origin, transcript abundance level (TPM) and the most likely allele the peptide would bind to as predicted by HLAthena using MSi (peptide sequence-only model) and MSiCE (cleavability and expression integrative model). **B**) Mirror plots for the MS spectra of two neoantigen sequences (LLHTELERFL and NSKKKWFLF) in the discovery experiment (top) and the corresponding synthetic peptide spectrum (bottom, red: y-ion, blue: b-ion, green: internal ion). **C**) Targeted MS experiment for mutated peptide YIHGRGWAL using a heavy peptide. After optimization, transitions of y6-and y7-ion were acquired of the heavy peptide alone (left, blank) and of the heavy peptide spiked into a peptidome sample of MEL6. Endogenous peptide detection is shown on the top, detection of the heavy peptide is shown at the bottom. **D**) CD137 signal of neoantigen specific TCRs in different target backgrounds (PMA/Ionomycin = positive control, Mel -tumor cell line, PBMCs from the corresponding patient, APCs pulsed with DMSO, mutant or wildtype peptide). Percent upregulation is shown for each panel.

Three of the detected neoantigens were predicted as strong binders to one of the patient’s respective alleles (*HLAthena* MSi and MSiCE, rank <0.5) and one was predicted as a weak binder (rank <2). For the two peptides identified in MEL 1 with predicted binding above the rank threshold (LLHTELERFL, MSiCE rank 4.07 and NSKKKWFLF, MSiCE rank 3.73), we confirmed that our identifications were correct using synthetic peptides (**Figure 5B**). The neoantigen YIHGRGWAL from NCEH1 detected in MEL 6 was also evaluated using targeted MS with heavy-labeled peptide. We observed the selected MS2 transitions for this peptide in an HLA peptide pool injection (six IPs, ∼300 ⨯10^6^ cells) but not in a blank injection (**Figure 5C**). Triplicate analysis across HLA IP samples from different cell culture batches confirmed the presentation of this neoantigen.

We then tested the reactivity of peptide-specific T cell receptor (TCR) clones against three of these MS-detected neoantigens using the CD137 signal as a marker for TCR transduction and activation. When cultivated with patient-derived melanoma cells that presented the antigens KRPTPTFHF, YIHGRGWAL, QLPDFVNQL at physiological levels as well as with antigen presenting cells (APC) pulsed with mutant peptides, TCRs were activated (**Figure 5D**, MEL track (red) and APC-Mut pep track). In contrast, incubation of peripheral blood mononuclear cells (PBMCs), or APCs pulsed with DMSO or WT peptides did not lead to TCR activation. In summary, our improved workflow enabled the detection of clinically relevant antigens that can initiate a tumor-specific T cell response.

## Discussion

To gain depth in the assessment of the HLA peptidome most studies have used billions of cells as input, numbers that typically cannot be obtained from clinical specimens, whether they are tumor specimens procured through standard biopsy or surgery, or even from cell line experiments where the number of obtainable cells is limited. Therefore, more sensitive approaches are needed to enable routine HLA peptidome analysis for clinical samples.

Here, we demonstrated the ability to achieve deep coverage of the HLA peptidome from cells and tissues using input amounts of as little as 100⨯10^6^ cells or 0.15g wet weight of tumor. Further increases in peptide yield relative to repeat injection of the same sample were obtained by fractionating the peptide specimen offline into 3 fractions using microscaled bRP on Stage-Tips. The use of SDB-XC material separated the peptidome in an allele-(and hence amino acid composition) dependent manner. By evaluating this method across a range of diverse HLA-I alleles in both mono-and multi-allelic settings, we observed that, although individual fractions reflected peptide characteristics obtained when binding and eluting under basic conditions, the total peptidome per sample was not biased to any particular allele. Offline bRP fractionation using other materials such as C18 was also feasible but may lead to the loss of the orthogonality to the on-line acidic RP separation afforded by the SDB-XC material. We noted that only 3 fractions were required per experiment, and thus measurement time remained comparable to our standard approach using 2 replicate injections.

Gas phase separation of ions using FAIMS increased the HLA-I eluted peptide yield compared to acquisition without FAIMS by reducing co-isolation of peptide precursors, and mitigated instances of: (i) one precursor would not be triggered, (ii) the ions would co-fragment and lead to mixed spectra without FAIMS, and (iii) removal of singly and highly charged contaminants introduced by the immunoprecipitation in one or the other CV. The usage of a combination of CVs (−50 and -70 on the Thermo FAIMSPro device) yielded the widest range of peptides. While singly charged ions are likely not acquired with these settings, the fraction of HLA-I eluted peptides lost was generally negligible compared to the overall gain in identifications. When foreseeing a sample composition lacking in basic residues and dominated by potentially nonpolar side chains likely to lead to a substantial proportion of singly charged peptide precursors (e.g., allele B*07:02), we propose adding a third experiment with a CV of -20 (**Supplemental Figure 3D**,**E**) ^17^. We expect similar benefits for use of ion mobility for HLA-I eluted peptides analysis with other types of instruments such as the TimsTOFpro from Bruker.

We also observed that variation in sample preparation, as well as data acquisition methods for the same sample, could affect the overall peptide yield and reveal unique peptides not found with more than one of the methods used. Maintaining optimal instrument performance is also an important consideration. Such variability in the identified peptide population due to sample preparation and data acquisition have been observed by others ^19,27,29,30^, but is not yet fully understood. Based on these findings, we developed a decision tree for different scenarios, dependent on the amount of available sample (**Figure 4E**). Specifically, if only material for a single acquisition is available, our current preferred method is bRP fractionation together with FAIMS-based acquisition, given our success in using this approach to detect neoantigens. If limiting instrument time is a concern, we still recommend using FAIMS while giving consideration to reducing the number of fractions or simply injecting the same sample twice using different CV sets for each acquisition ^20^. In our experience, repeat technical injections provided the lowest yield of HLA-I eluted peptides; thus, we recommend varying acquisitions by using bRP fractionation and FAIMS alone and in combination in cases where sufficient sample is available.

The increase in sensitivity for detection of HLA-I eluted peptides obtained by combining bRP fractionation and FAIMS acquisition enabled the identification of neoantigens from tumor-derived cell line samples that were not found in single-shot experiments. We confirmed that MS-detected neoantigens can induce immune responses and are instrumental for the identification of cognate antigens of TCRs as well as selection of potential vaccine targets (Oliveira et al., in revision).

Future improvements to these sample preparation and data acquisition methods may include but are not limited to: (i) the use of automation for HLA peptide enrichment ^31^, (ii) online LC-columns of smaller inner diameter ^32^ or (iii) shorter gradients in combination with FAIMS acquisition methods ^33^. Chemical labeling has also been shown to increase the number of peptide identifications, but may potentially change the overall allele representation in the sample ^19^.

In conclusion, offline separation and gas phase fractionation increased the yield of HLA-I presented peptides analyzed by LC-MS/MS and allowed for detection of low abundant neoantigens. We foresee that similar observations are true for HLA class II peptides and that these acquisition approaches will benefit antigen discovery not only in cancer, but also in other fields, including autoimmune and infectious diseases.

## Acknowledgments

The authors thank Sebastian Vaca and Pierre M. Jean-Beltran for computational assistance. The authors thank Scott Goulding for experimental discussions. S.K. was supported by the Cancer Research Institute as a Hearst Foundation fellow. This work was made possible by a grant from BroadIgnite at the Broad Institute of MIT and Harvard. D.B.K. was supported by the Emerson Collective. This work was also supported in part by grants from the National Cancer Institute (NCI) Clinical Proteomic Tumor Analysis Consortium grants NIH/NCI U24-CA210986 and NIH/NCI U01 CA214125 (to S.A.C). This work was supported in part from grants from the National Institutes of Health (NCI-1R01CA155010; NCI-U24CA224331 [to C.J.W.]; NIH/NCI R21 CA216772-01A1, NCI-R01 CA229261 [to P.A.O] and NCI-SPORE-2P50CA101942-11A1 [to D.B.K], NCI-SPORE P50CA101942-15)[to D.A.B.], DOD CDMRP (KC190128) [to D.A.B] and a Team Science Award from the Melanoma Research Alliance (C.J.W., P.A.O.). G.O. was supported by the American Italian Cancer Foundation fellowship. This work was further supported by The G. Harold and Leila Y. Mathers Foundation

## Supplemental Figures

**Supplemental Figure 1.**
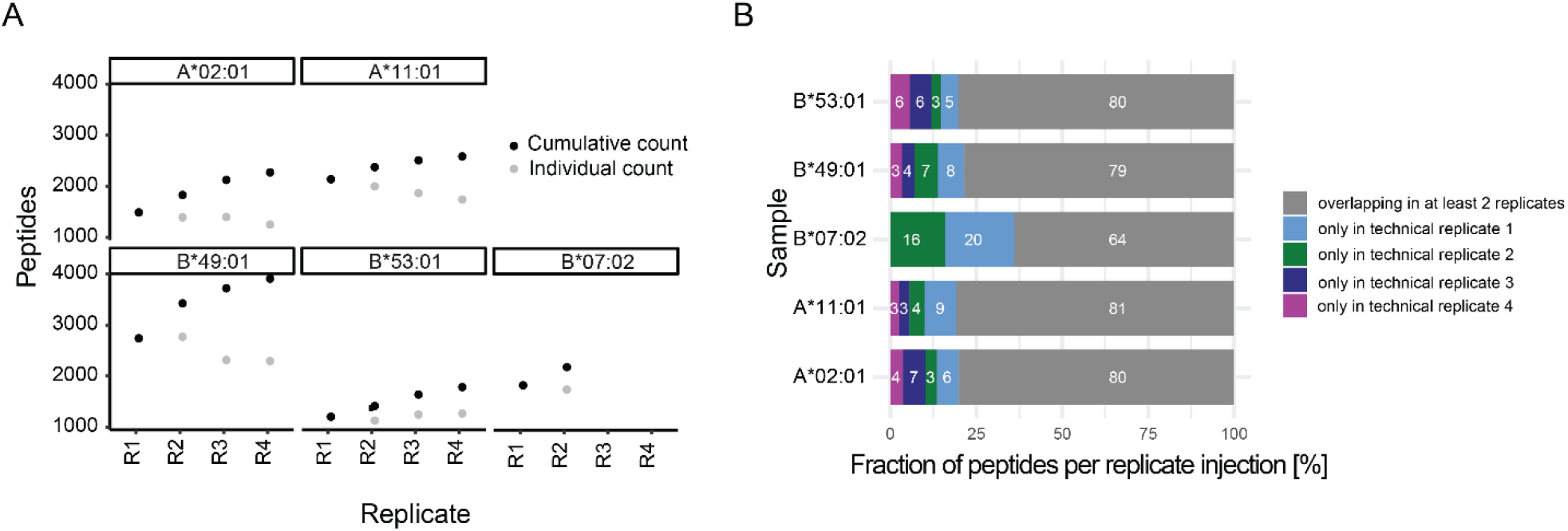
**A**) Cumulative increase of peptide identifications in repeat injections (black dots) across multiple alleles as profiled in [8]. Grey dots indicate individual peptide yield per injection. **B**) Barchart showing the overlap in identifications in the alleles from **A**. Grey fraction are peptides identified in more than one injection, increase by each additional injection is marked in a different color.

**Supplemental Figure 2.**
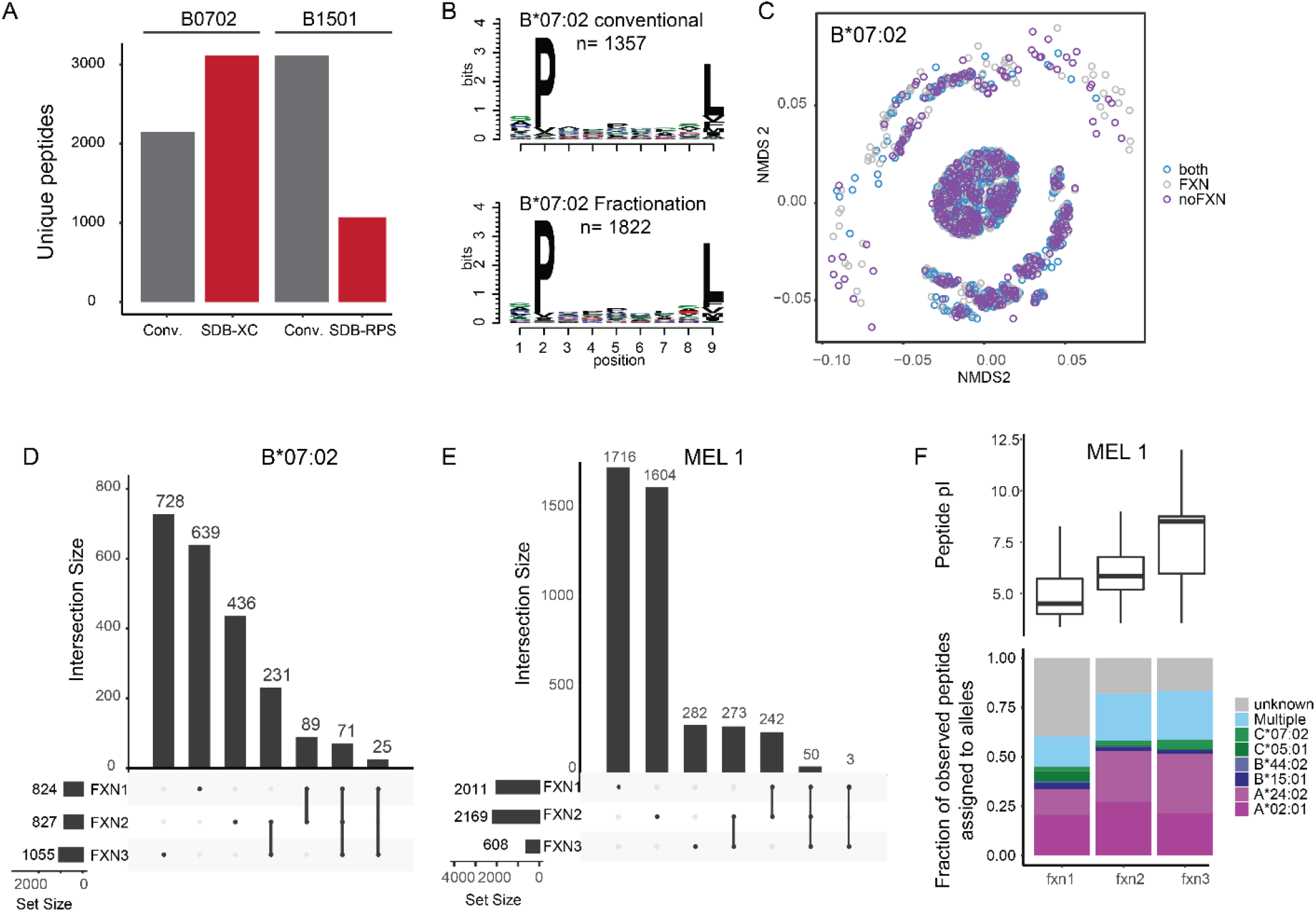
**A**) Number of unique peptide identifications in fractionated sample preparations of HLA-B*07:02 on SDB-XC material and HLA-B*15:01 peptides on SDB-RPS material compared to unique identifications in two technical replicate injections without fractionation of the same allele. **B**) 9mer motif of additional B*07:02 binders identified using no fractionation (conventional, top) or fractionation (bottom) approaches. **C**) NMDS plot of B*07:02 binders identified in samples using conventional (blue), fractionated (purple) and both (grey) methods. **D)** Peptide spectrum matches (PSMs) identified across the three fractions of the B*07:02 peptidome. Horizonal bars, left bottom, indicate the number of total PSMs per fraction, dots and lines indicate whether vertical bar chart PSMs are identified in only one (single dot) or multiple fractions (dots with lines). **E**) Same as **D** but for MEL1. **F**) Allele assignment of peptides identified in the individual fractions of MEL 1(left, HLAthena rank <0.5). The boxplot shows peptide pI of peptides per fraction.

**Supplemental Figure 3.**
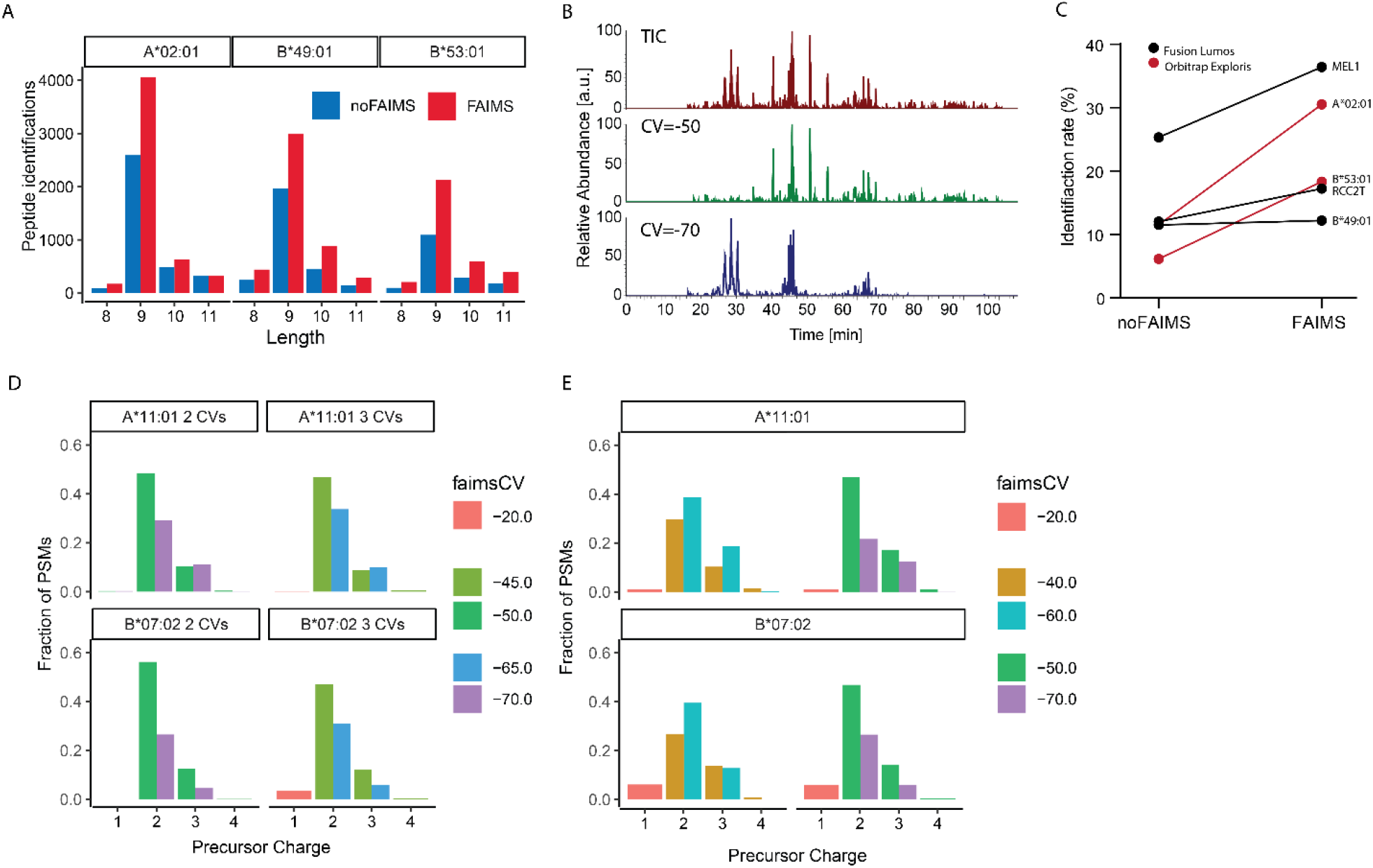
**A**) Length distribution for A*02:01, B*49:01, and B*53:01 peptides identified with and without FAIMS. **B**) Total Ion Chromatogram (TIC) of an HLA-peptide acquisition in total (top) and separated by CV (middle CV -50, bottom CV -70). **C**) Identification rate improves when using FAIMS. **D**) Charge state distribution comparing two acquisitions of A*11:01 and B*07:02 using 2CVs (−50|-70) or 3 CVs (−20|-45|-65). **E**) Comparison of two 3 CV methods (−20|-40|-60 or -20|-50|-70) and their impact on charge states on two alleles A*11:01 and B*07:02.

**Supplemental Figure 4.**
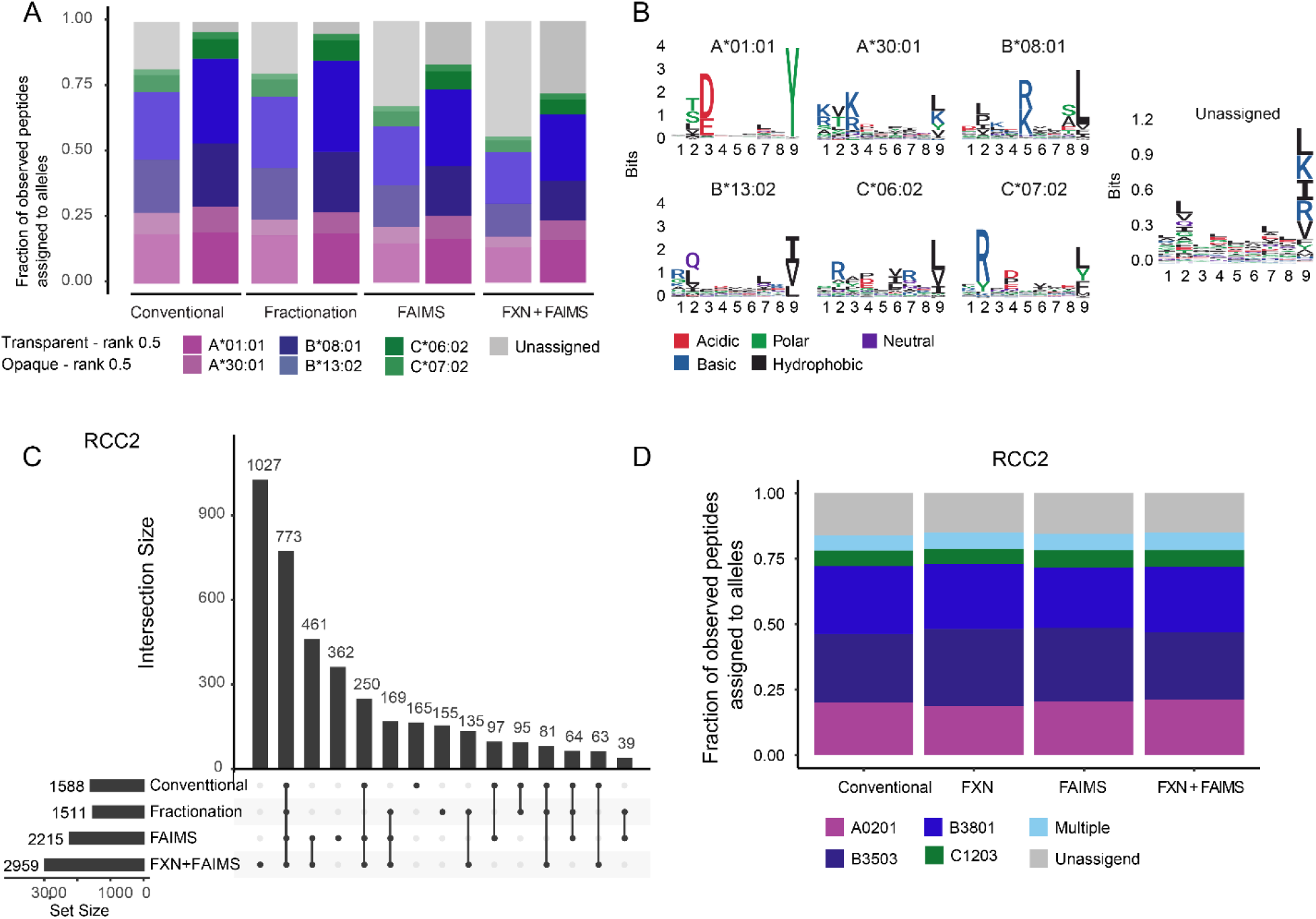
**A**) Allele assignment for RCC4T at rank <0.5 (transparent bars) and rank <2 (colored bars). **B**) Logo plots for peptides assigned to each allele across all experiments and logo plot for unassigned peptides in the Fractionation + FAIMS run. **C**) Peptides identified across all acquisitions of RCC2T. Left bar charts indicate the number of total peptides per experiment, dots and lines indicate whether vertical bar chart peptides are identified in only one (single dot) or multiple fractions (dots with lines). **D**) Allele assignment for RCC2T.

